# Responses to novel food resources in the extinct-in-the-wild ‘alalā

**DOI:** 10.64898/2026.07.11.737884

**Authors:** Averill Cantwell, Bryce Masuda, Alison L. Greggor

## Abstract

One challenge of reintroduction programs is ensuring that captive individuals are prepared for life in the wild, including having the ability to forage on wild resources. Foraging proficiency can be especially challenging to foster in species with a wide dietary breadth or reliance on ephemerally available resources. A relatively unexplored obstacle to release readiness may be an animal’s hesitancy to approach or consume unfamiliar foods, i.e. their neophobia and dietary conservatism. In this study, we examined the responses of a conservation breeding population of the extinct-in-the-wild ‘alalā (*Corvus hawaiiensis*), a dietary generalist, to novel and familiar native and non-native fruits. Birds were similarly willing to contact both novel and familiar fruits, suggesting little aversion to the presence and appearance of novel food items. However, birds showed a clear preference for contacting non-native fruits over native fruits, regardless of familiarity. Additionally, while birds were willing to contact novel items, they were less likely to consume novel native fruits, suggesting that dietary conservatism may limit incorporation of these resources in the wild. Finally, juvenile birds were less likely to approach the testing setup and contact items, suggesting that neophobia may constrain their exploration of novel environments and resources. We discuss the implications of ‘alalā preferences for non-native foods and reluctance to consume novel native items for pre-release preparations and the goal of wild species recovery.

## Introduction

Reintroduction programs are critical tools in the recovery of endangered species. When reintroductions rely on captive-bred animals, a major challenge is ensuring these individuals are adequately prepared for life in the wild, including demonstrating the ability to efficiently identify and consume wild resources (Greggor et al., 2024; Shier, 2016). Captive-bred animals often forage less efficiently than their wild counterparts, which can reduce post-release survival and reproductive success (Beck et al., 1994; Brown & Laland, 2001; Ellis et al., 2000; Kleiman, 1989; Mathews et al., 2005; Owen-Smith & Festa-Bianchet, 2003; Stoinski et al., 2003; Stoinski & Beck, 2004). Multiple factors may contribute to these deficits, such as the time required to learn the spatial and temporal distribution of resources (Owen-Smith & Festa-Bianchet, 2003) and reduced foraging effort by captive-bred individuals after release to the wild (Stoinski et al., 2003; Stoinski & Beck, 2004). Another key factor may be how animals respond to novel food resources.

Neophobia–the avoidance of novel stimuli–is an important influence on foraging behavior. In many contexts, neophobia is adaptive, protecting an animal against ingesting harmful or dangerous items (Mettke-Hofmann et al., 2002; Mettke-Hofmann, 2014). From a conservation management perspective, however, neophobia can pose an issue if captive individuals avoid nutritionally appropriate but unfamiliar resources following release into the wild. While it is possible to expose some specialist species to their complete diet in human care, this is often infeasible for generalist species that consume a broad and seasonally variable range of resources.

Even when an initial neophobic response is overcome, many animals continue to exhibit dietary conservatism–reluctance to consume a novel food, even if they are willing to approach or contact the item (Greenberg & Mettke-Hofmann, 2001; Marples & Kelly, 1999; Marples et al., 2007). While overcoming neophobia is a necessary step in addressing dietary conservatism, these are thought to be distinct processes, with dietary conservatism requiring a longer, slower process before an animal accepts a new food item into its diet (Marples & Kelly, 1999). Understanding and addressing these factors is a crucial component of reintroduction programs. For example,pre-release exposure to wild foods boosted post-release survival in multiple studies of translocated birds (Roberts & Luther, 2023), though the specific role of reduced neophobia or dietary conservatism in these changes was not explored. Overall, an understanding of the impact of neophobia and dietary conservatism on pre-release preparations remains limited to relatively few species.

Here, we examine the potential influence of food neophobia and dietary conservatism on foraging decisions in the extinct-in-the-wild ‘alalā (*Corvus hawaiiensis*) (IUCN, 2025), also known as the Hawaiian crow, which is the focus of an ongoing reintroduction program. ‘Alalā went extinct in the wild in 2002, and current recovery efforts rely entirely on individuals bred and reared in human care (U.S. Fish and Wildlife Service, 2009). Historically, wild ‘alalā consumed a broad diet that included native fruits, invertebrates, nectar, plant parts, eggs, nestlings, and small vertebrates (Sakai et al., 1986; Sakai & Carpenter, 1990). In particular, their consumption of a high diversity of plant species is thought to have made ‘alalā critical seed dispersers within the Hawaiian forest ecosystem (Culliney et al., 2012). While wild birds occasionally foraged on non-native foods such as bananas and gourds, these items are believed to have been a small proportion of their diet, particularly given how poorly birds fared in human-dominated landscapes (Baldwin, 1969; Banko et al., 2020). The diet provided to the captive population aims to replicate the historical diet nutritionally, but it is currently infeasible to supply the full diversity and quantity of fruits, insects, and other items consumed by wild birds. Consequently, understanding how captive ‘alalā respond to novel food resources is important for predicting potential barriers to post-release foraging success.

Neophobia is often less intense in dietary generalists, who exploit a wide range of food resources (Clarke & Lindburg, 1993; Glickman & Sroges, 1966; Greenberg & Mettke-Hofmann, 2001; Webster & Lefebvre, 2000). However, despite broad diets, many corvid species nonetheless exhibit high levels of neophobia, including towards novel foods (Brown & Jones, 2016; Greggor et al., 2016a; Greggor et al., 2016b; Miller et al., 2022). Previous work has shown that ‘alalā display strong neophobia toward novel objects, especially as juveniles (Greggor et al., 2020), and at levels exceeding those of related species (Miller et al., 2022). Whether this object neophobia extends to high levels of food neophobia is not yet clear, as object and food neophobia are thought to reflect distinct motivations (Greggor et al., 2015). The role of dietary conservatism has also not yet been explored. If ‘alalā exhibit strong food neophobia or dietary conservatism, especially towards native species they will encounter post-release, this may impact how effectively ‘alalā forage in the wild.

The goal of the present study is to assess how ‘alalā in a conservation breeding and reintroduction program respond to novel and familiar food items, including both native and non-native fruits, prior to release. The native fruits selected reflect those ‘alalā are likely to encounter in the wild. We also investigate whether juveniles and adults differ in their willingness to approach, contact, and consume novel foods, as age-related differences could have important implications for pre-release preparations or age recommendations for release. Finally, while past work in ‘alalā did not detect sex differences in object neophobia, we include sex as females of some species exhibit greater neophobia than males in response to novel objects (e.g., Ensminger & Westneat, 2012) and environments (e.g., King et al., 2013).

## Methods

### Housing

Birds were housed at the Keauhou Bird Conservation Center, as part of a wider reintroduction program for the species. All birds were housed in multi-chamber aviaries with access to ambient light, temperature, and precipitation, and had ad libitum access to food and water throughout the duration of the trial. Testing occurred in birds’ home enclosures. Birds were housed with a mate, a small group of conspecifics, or in a single chamber with visual and acoustic access to conspecific neighbors. Aviaries contained native wood perching and browse, and all were part of an enrichment program that aimed to offer opportunities to display species-typical behavior (see Greggor et al., 2018 for details).

### Diet

‘Alalā are omnivorous, naturally foraging on fruits, insects, flower nectar, plant parts, and small passerine eggs and nestlings (Banko et al., 2002; Sakai et al., 1986). The daily diet provided to this population aimed to replicate what was believed to be the nutritional composition of their wild diet. This included papaya, apple, frozen mixed vegetables, cooked chicken egg, frozen mice and nutritionally balanced pellets (Zupreem Monkey biscuits and Mazuri softbill pellets). Birds were also given opportunities to interact with and consume a rotation of native and non-native fruits, along with other enrichment items. To determine which fruit species were familiar versus novel, we cataloged four years of enrichment records (over 2,000 records) to confirm what had previously been provided. A total of 49 plant types had been given during this time, including 14 native fruits. The most common native fruit type was pūkiawe (*Leptecophylla tameiameiae*) (N = 109). The most common non-native fruits were watermelon (*Citrullus lanatus*) (N = 39) and grapes (*Vitis vinifera*) (N = 33). The novel fruits chosen for this study had not appeared in enrichment records during that time, nor could any current staff member recall them being fed in the 10 years prior to record keeping. We expected ‘alalā to be capable of remembering the familiar fruits offered to them, as they historically foraged on dozens of species that are only available seasonally (Banko et al., 2002).

Additionally, evidence from other corvid species suggests strong long-term memory in other contexts (at least 10 months (Bogale et al., 2012); at least 4 years (Blum et al., 2020)) as well as episodic memory abilities associated with food caching (Grodzinski & Clayton, 2010).

### Ethics statement

This study was approved by San Diego Zoo Wildlife Alliance’s IACUC committee (No. 16–009). The conservation breeding of ‘alalā was covered by: U.S. Fish and Wildlife Service Native Endangered Species Recovery Permit TE060179-5, State of Hawaii Protected Wildlife Permit WL19-16. We collected native fruits with permission from the State of Hawaii Department of Land and a Natural Resources Access and Forest Reserve Special Use Permit.

### Subjects

We tested the responses of 54 birds (23 females and 31 males, age 4 months to 21 years, mean: 7, SD: 5) from August to November 2018. At the time of the study, the Keauhou Bird Conservation Center housed 82 ‘alalā. The testing effort included as many birds as possible given aviary layout constraints and veterinary or husbandry requirements. We classified juveniles as birds under 3 years old, and all other individuals as adults, based on age at sexual maturity and in line with other neophobia analyses (Greggor et al., 2020; Banko et al., 2002), although breeding can occur at 2 years (Barrett et al., 2024). This resulted in age classes of 10 juveniles and 44 adults.

### Procedure

We tested birds over a series of four conditions. In each trial, a control fruit and a test fruit were placed inside the bird’s aviary on the right and left sides of a familiar platform or ledge (Figure 1). The location for testing was consistent for all trials within each aviary, and the control and test fruits were counterbalanced right and left across trials. We included a control fruit– papaya, a highly familiar food included in the daily diet of all birds–to verify the bird’s motivation and willingness to approach and contact a food item in the test setup. This allowed us to disentangle hesitancy to approach potential novelty from a lack of interest in food. There were four categories of test fruit: 1) novel non-native fruit (dragonfruit, *Selenicereus undatus*), 2) novel native fruit (kolea, *Myrsine lessertiana* or kūkaenēnē, *Coprosma ernodeoides*), 3) familiar non-native fruit (grapes, *Vitis vinifera* or watermelon, *Citrullus lanatus*), and 4) familiar native fruit (pūkiawe, *Leptecophylla tameiameiae*). See Figure 2 for images of each native fruit type.

**Figure 1.**
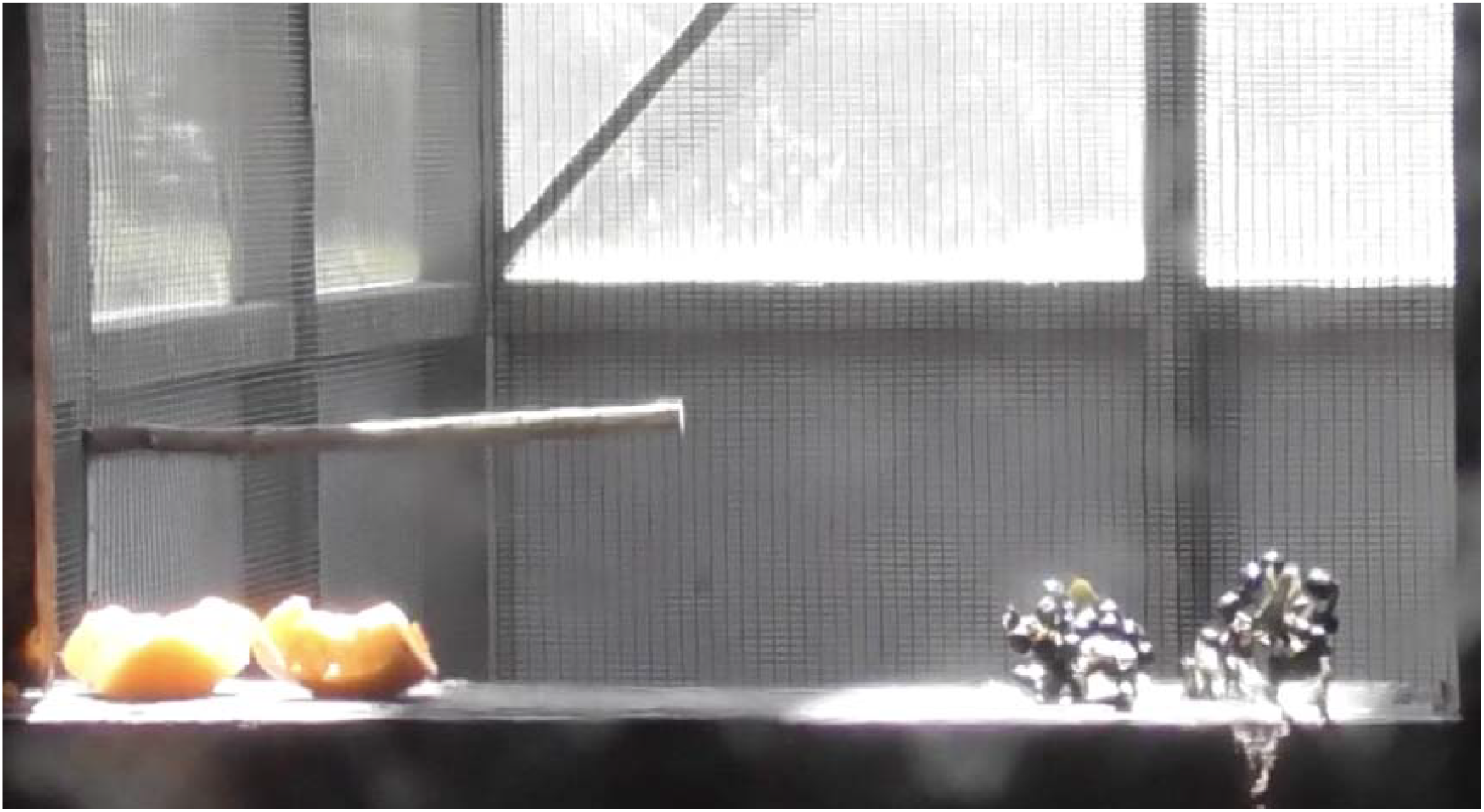
Testing setup. The control fruit (papaya, left) and test fruit (kolea, right) were placed on either side of a ledge within the bird’s aviary.

**Figure 2.**
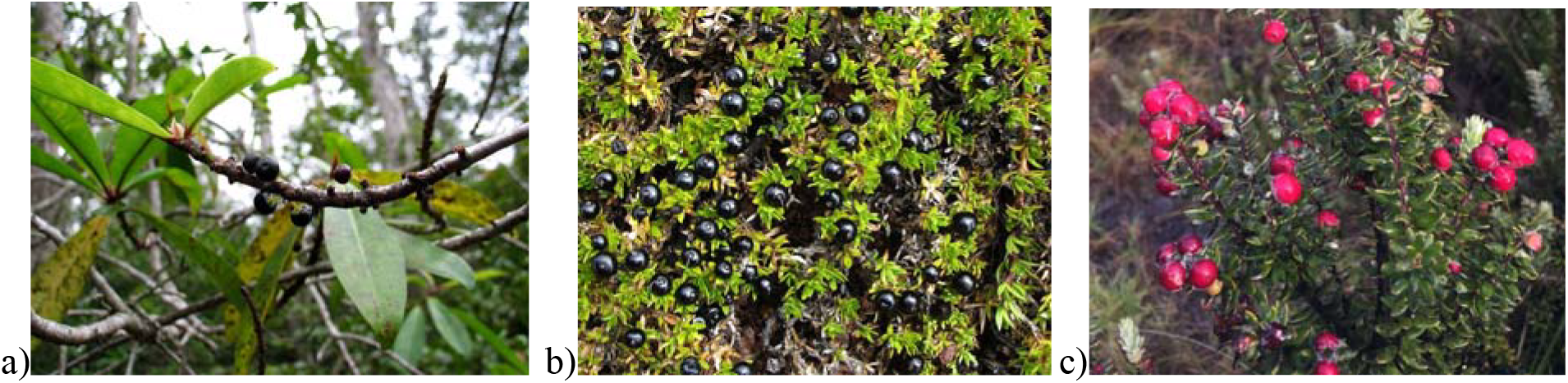
Native fruit types. (a) Kolea (*Myrsine lessertiana*), (b) Kūkaenēnē (*Coprosma ernodeoides*); photos by David Eickhoff, CC BY-SA, (c) Pūkiawe (*Leptecophylla tameiameiae*); photo by Alison Greggor.

We conducted trials on separate days with a break of at least one day in between and randomized trial order. 29 birds completed trials with all four categories of test fruit, 17 birds completed trials with three categories, 5 with two categories, and 3 with only one category. Within the novel native fruit category, 35 birds completed two trials (one with kolea and one with kūkaenēnē) to help determine if there were differences between native fruit exemplars. Birds who only completed a subset of trial types did so due to unrelated testing constraints, such as veterinary or husbandry needs. In some trials, it was not possible to conclusively identify the bird in view; these trials were excluded. We left food items inside the aviary for approximately 30 minutes and responses were recorded on a freestanding camcorder, placed outside the aviary with a view to the test location through an observation window. No humans were present apart from setup and breakdown of the trial.

### Coding and data analysis

All trials were recorded on video. A coder identified each bird in the video by their unique combination of colored leg bands and coded (1) whether an individual approached the ledge where the fruits were placed, (2) latency to contact the control and/or test fruit, and (3) whether they were observed consuming the control and/or test fruit. A separate observer re-scored ~20% of trials (25 of 121 videos; 47 of 215 rows of data), with high reliability for (1) approach (κ = 1), (2) latency to contact the control (*r* > 0.99) and/or test (*r* > 0.99) fruit, and (3) consumption of control (κ = 1) and/or test fruit (κ = .91).

We analyzed data in R 4.4.1 (R Core Team, 2024). First, to assess willingness to approach the testing setup we built GLMMs with a binomial error distribution and a logit link function using the *glmer* function from the *lme4* package (Bates et al., 2015). To assess latency to contact different types of fruit, we used a Cox proportional hazards regression model using the *coxph* function from the *survival* package (Therneau, 2015). Finally, to assess willingness to consume different fruit types, we again used GLMMs with a binomial error distribution. In all analyses we included *subject* as a random effect, and considered *sex, age class* (juvenile or adult), *fruit novelty* (familiar or novel), *fruit category* (native or non-native), and the interaction between *novelty* and *category*. We used the *MuMIn* package to run model selection based on AICc values. If multiple models had a ΔAICc within 4 of the top model, we averaged models (Burnham & Anderson, 2002; Grueber et al., 2011). We interpreted the influence of averaged variables based on their relative importance scores (RI), giving more weight to higher scores.

## Results

We first assessed overall willingness to approach the testing area. Rates of approach were similar when comparing native (mean=0.90 ± se=0.03) and non-native fruit trials (0.84 ± 0.04) (Fisher’s exact test: p = 0.23), as well as familiar (0.84 ± 0.04) and novel (0.90 ± 0.03) fruit types (p = 0.23). However, rates of approach differed by age class: adults approached the testing setup at high rates (0.93 ± 0.02) compared to juvenile birds (0.59 ± 0.08) (p < 0.0001, Figure 3). GLMMs confirmed this result: only *age class* had a high relative importance score in our averaged model (N = 215 trials, RI□=□1.00), indicating that adult birds were more likely to approach the testing setup (*β*□=□1.98, *z*□=□3.28; Table 1, 2).

**Table 1.** Predictors included in averaged models.

| (Intercept) | Age Class | Fruit Category | Fruit Novelty | Sex | Category* Novelty | df | logLik | AICc | delta | weight |
| --- | --- | --- | --- | --- | --- | --- | --- | --- | --- | --- |
| 1.90 | + |  |  |  |  | 3 | -69.49 | 145.09 | 0.00 | 0.31 |
| 2.12 | + | + |  |  |  | 4 | -69.04 | 146.27 | 1.18 | 0.17 |
| 1.66 | + |  | + |  |  | 4 | -69.07 | 146.32 | 1.23 | 0.17 |
| 1.67 | + |  |  | + |  | 4 | -69.28 | 146.76 | 1.66 | 0.13 |
| 1.88 | + | + | + |  |  | 5 | -68.71 | 147.71 | 2.62 | 0.08 |
| 1.88 | + | + |  | + |  | 5 | -68.82 | 147.93 | 2.84 | 0.07 |
| 1.44 | + |  | + | + |  | 5 | -68.88 | 148.04 | 2.95 | 0.07 |

**Table 2.** Output of averaged GLMM predicting likelihood of approaching the testing setup.

|  | Estimate | Std. Error | Adjusted SE | z value | Pr(>\|z\|) | RI |
| --- | --- | --- | --- | --- | --- | --- |
| (Intercept) | 1.83 | 0.56 | 0.56 | 3.25 | <0.01 |  |
| **AgeClassAdult** | **1.98** | **0.60** | **0.60** | **3.28** | **<0.01** | **1.00** |
| **FruitCategoryNonNative** | -0.15 | 0.35 | 0.35 | 0.42 | 0.68 | 0.33 |
| **FruitNoveltyNovel** | 0.14 | 0.34 | 0.34 | 0.40 | 0.69 | 0.32 |
| **SexMale** | 0.12 | 0.42 | 0.42 | 0.29 | 0.77 | 0.28 |

**Table 3.** Predictors included in averaged models of latency to contact test fruit.

| (Intercept) | Age Class | Fruit Category | Fruit Novelty | Sex | Category* Novelty | df | logLik | AICc | delta | weight |
| --- | --- | --- | --- | --- | --- | --- | --- | --- | --- | --- |
| + | + | + |  |  |  | 2 | -628.80 | 1261.7 | 0.00 | 0.43 |
| + | + | + | + |  |  | 3 | -628.70 | 1263.6 | 1.88 | 0.17 |
| + | + | + |  | + |  | 3 | -628.73 | 1263.6 | 1.95 | 0.16 |
| + | + | + | + |  | + | 4 | -628.05 | 1264.4 | 2.71 | 0.11 |
| + | + | + | + | + |  | 4 | -628.62 | 1265.5 | 3.84 | 0.06 |
| + |  | + |  |  |  | 1 | -631.81 | 1265.6 | 3.95 | 0.06 |

**Figure 3.**
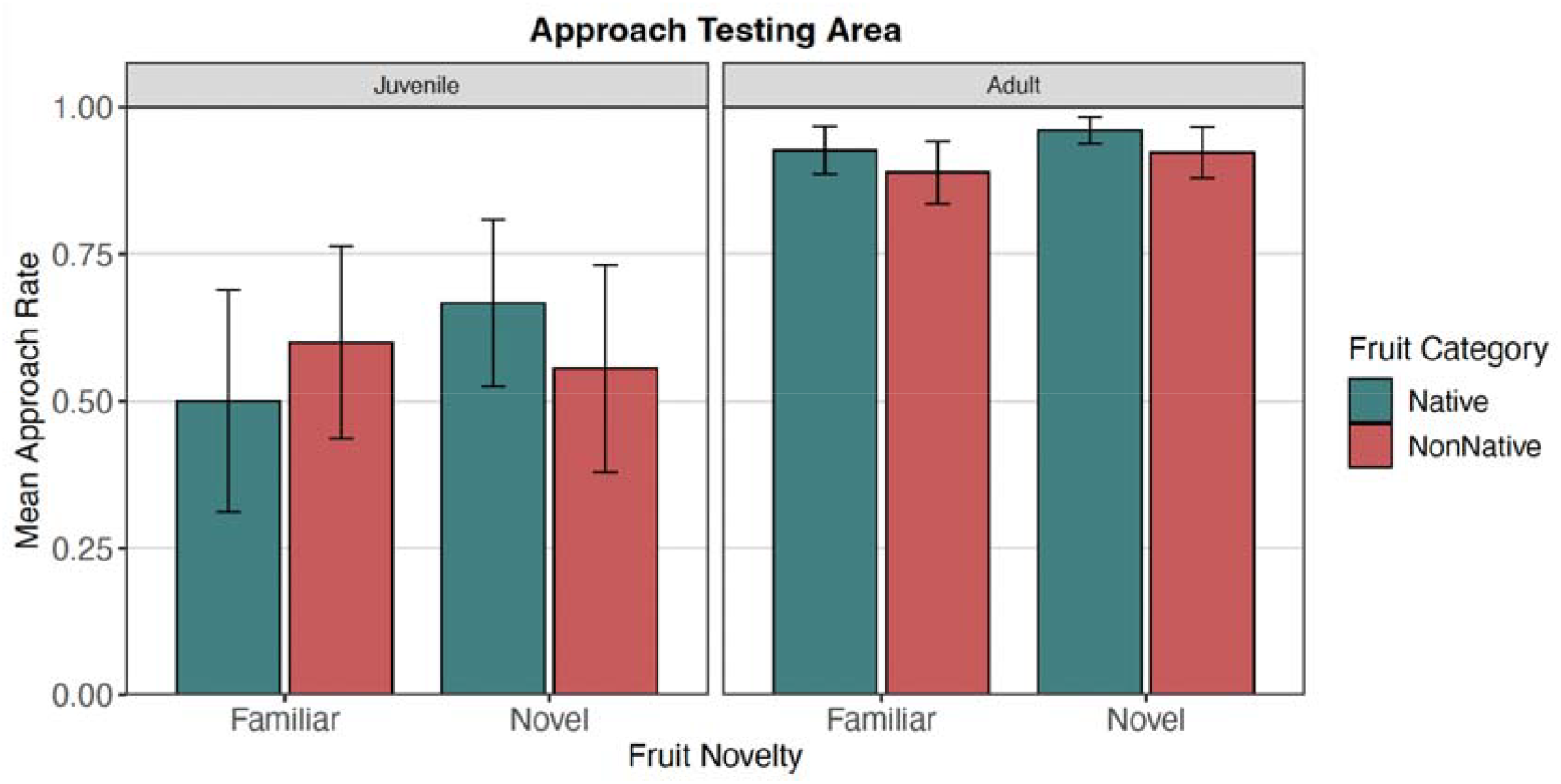
Willingness to approach the testing area by age, fruit category, and fruit novelty (mean ± SE).

We next analyzed latency to contact the test fruit among birds who approached the setup (N = 185 trials). Similar to the previous result, juvenile birds who approached the testing setup were less likely to contact the test fruit compared to adults (*β*□=□0.42, *z*□=□2.52, RI = 0.94; Table 4, Figure 4a). In addition, birds were more likely to contact non-native fruits compared to native fruits (*β*□=□0.63, *z*□=□2.91, RI = 1.00; Table 4, Figure 4c). We also compared these results to a model predicting latency to contact the control fruit (papaya) (see Supplemental Materials). In contrast to the test fruit, we did not find a difference between juvenile and adult birds’ likelihood of contacting the control fruit. We also found that birds were more likely to contact the control fruit when it was paired with a native fruit compared to a non-native fruit, in line with the reduced contact of native fruits found above, but found no difference based on novelty.

**Table 4.** Output of averaged Cox proportional hazards model testing factors influencing the likelihood of contacting the test fruit.

|  | Estimate | Std. Error | z value | Pr(>\|z\|) | RI |
| --- | --- | --- | --- | --- | --- |
| **AgeClassAdult** | **0.42** | **0.17** | **2.52** | **0.01** | **0.94** |
| **FruitCategoryNonNative** | **0.63** | **0.22** | **2.91** | **0.00** | **1.00** |
| **FruitNoveltyNovel** | 0.05 | 0.13 | 0.37 | 0.71 | 0.34 |
| **SexMale** | 0.02 | 0.10 | 0.15 | 0.88 | 0.23 |
| **Category\*Novelty** | -0.05 | 0.17 | 0.27 | 0.79 | 0.11 |

**Table 5.** Predictors included in averaged models.

| (Intercept) | Age Class | Fruit Category | Fruit Novelty | Sex | Category* Novelty | df | logLik | AICc | delta | weight |
| --- | --- | --- | --- | --- | --- | --- | --- | --- | --- | --- |
| -0.77 |  | + | + |  | + | 5 | -84.73 | 179.79 | 0.00 | 0.42 |
| -0.99 | + | + | + |  | + | 6 | -84.34 | 181.15 | 1.36 | 0.21 |
| -0.79 |  | + | + | + | + | 6 | -84.73 | 181.92 | 2.13 | 0.15 |
| -1.01 | + | + | + | + | + | 7 | -84.34 | 183.31 | 3.51 | 0.07 |

**Figure 4.**
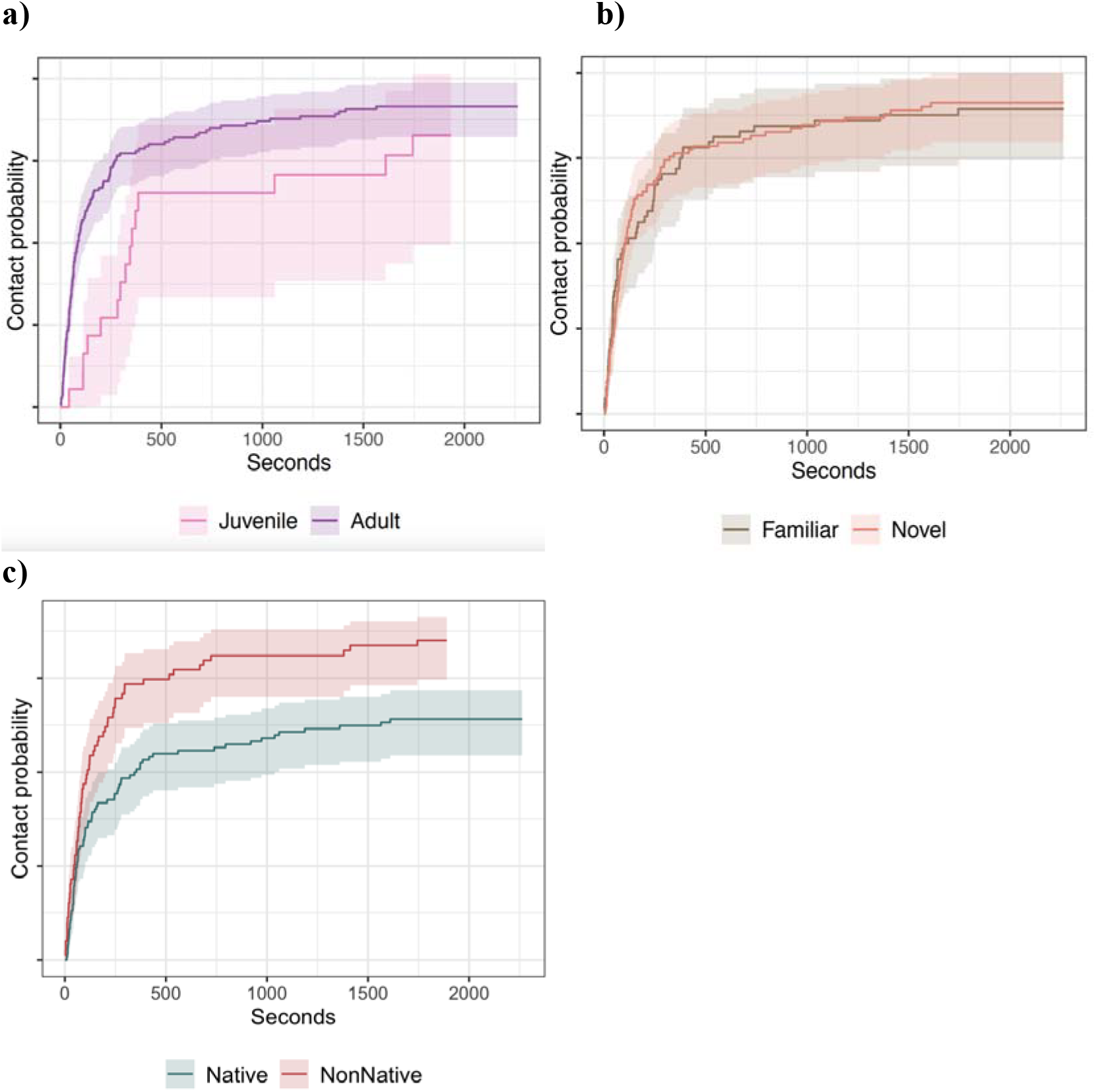
Probability of contacting the test fruit by age class, fruit novelty, and fruit category. (Survival curves estimated from models of isolated variables.)

Finally, we tested how these factors influenced whether the bird was observed consuming the test fruit. Overall, rates of consumption were low: only 26 birds were observed eating the test fruit in at least one trial. We again restricted our sample to birds who approached the testing setup. If a bird approached the testing setup, they ate the test fruit in only 19% of trials, compared to eating the control fruit (papaya) in 54% of approach trials.

We found an interaction between fruit category and fruit novelty in the likelihood that a bird was observed consuming the test fruit (N = 185 trials, *β*□=□2.26, *z*□=□2.79, RI = 1.00; Table 6). Birds were less likely to consume the novel native fruit compared to the familiar native fruit, while novelty did not have a significant effect on patterns of consumption for the non-native fruits (Figure 5). We additionally checked these results using only individuals who were observed eating at least one fruit (control, test, or both), to ensure this result was not simply driven by a lack of interest in food during the trial; these analyses yielded the same pattern (see Supplemental Materials).

**Table 6.** Output of averaged GLMM predicting likelihood of eating the test fruit.

|  | Estimate | Std. Error | Adjusted SE | **z**<br>value | Pr(>\|z\|) | RI |
| --- | --- | --- | --- | --- | --- | --- |
| (Intercept) | -0.85 | 0.43 | 0.43 | 1.98 | 0.05 |  |
| **FruitCategoryNonNative** | -0.69 | 0.55 | 0.55 | 1.25 | 0.21 | 1.00 |
| **FruitNoveltyNovel** | -1.75 | 0.56 | 0.56 | 3.11 | 0.00 | 1.00 |
| **CategoryNonNative\***<br>**NoveltyNovel** | **2.26** | **0.80** | **0.81** | **2.79** | **0.01** | **1.00** |
| **AgeClassAdult** | 0.14 | 0.34 | 0.34 | 0.40 | 0.69 | 0.34 |
| **SexMale** | 0.01 | 0.21 | 0.21 | 0.04 | 0.97 | 0.26 |

**Figure 5.**
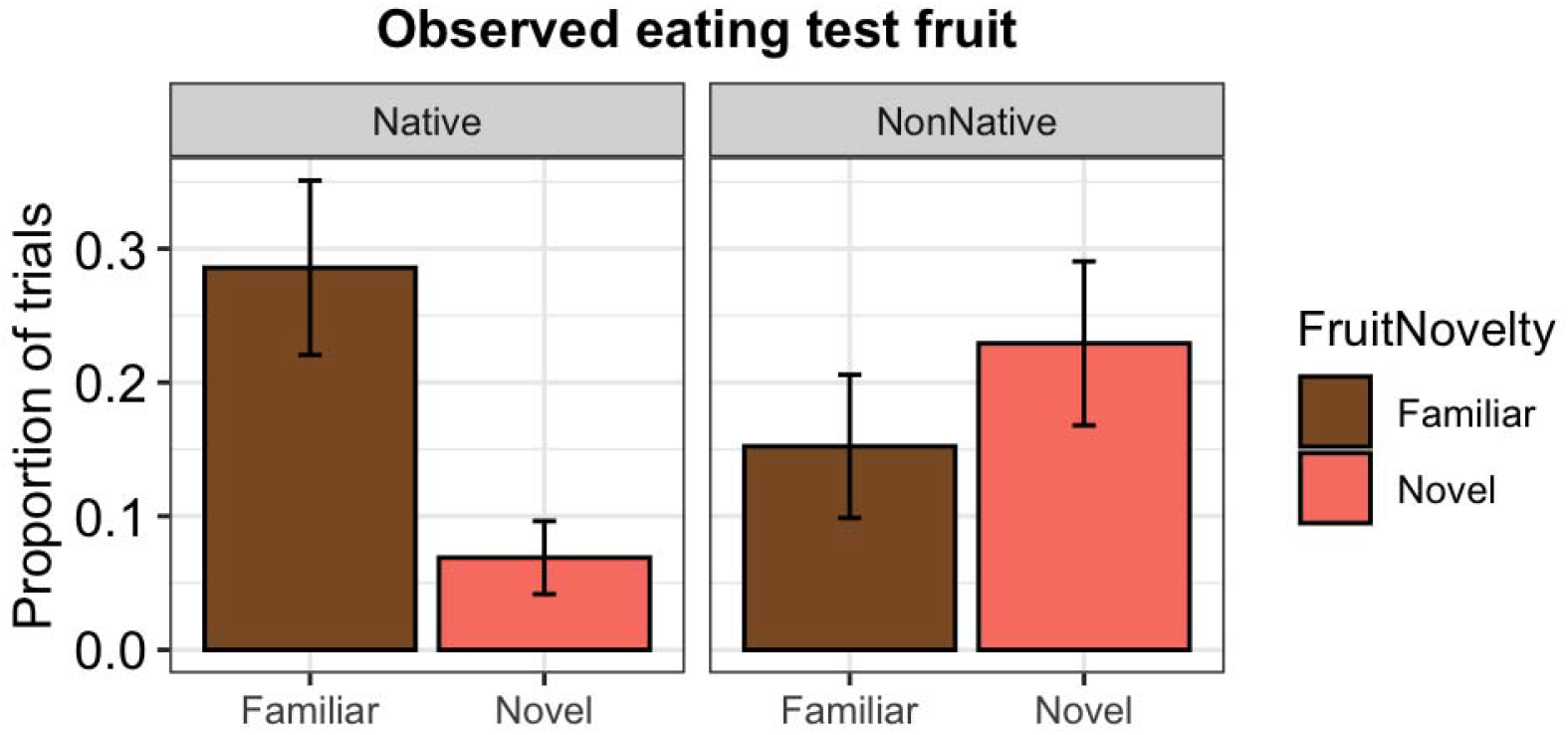
Patterns of test fruit consumption by novelty and category.

## Discussion

We investigated how the extinct-in-the-wild ‘alalā responds to food resources in order to better inform release preparation and management. Our results highlight several areas that may present challenges for ensuring captive ‘alalā are prepared to consume a diverse native diet after release. First, consistent with prior work showing greater object neophobia in juvenile ‘alalā (Greggor et al., 2020), juvenile birds were less likely to approach the testing setup, and, even when they did approach, less likely to contact the test fruit. Second, birds were generally less likely to contact native fruits, regardless of novelty. However, novelty did play a role in patterns of native fruit consumption: ‘alalā were less likely to eat novel native fruits compared to familiar native fruits, a difference not seen with non-native fruits. This preference for non-native resources and dietary wariness toward novel native foods may have important implications for the conservation management of this species.

Differences between age classes in how ‘alalā approach and contact food types are important considerations when selecting and preparing individuals for release. Although research on the effects of age on post-release outcomes is still a growing field, a recent review of bird translocations suggests that younger individuals often achieve better outcomes than adults (Miller et al., 2024). This may partly reflect the negative effects of prolonged captivity in older individuals, including stronger neophobic responses in captive-reared animals (Crane & Ferrari, 2017). Even in wild populations, neophobia is typically greater in adult animals compared to juveniles (reviewed in Greenberg, 2003). However, our results add to growing evidence that ‘alalā exhibit the opposite pattern, with juveniles exhibiting greater neophobia than adults (see also Greggor et al., 2020). Notably, juveniles did not appear to be food neophobic per se, but were less likely to approach and contact the novel testing setup, a pattern suggestive of object neophobia. This result could have mixed implications for post-release outcomes. Depending on the context, heightened object neophobia may protect juvenile birds from novel threats; however, if juvenile ‘alalā are less willing to approach unfamiliar resources in the wild, they may not adopt a species-appropriate diet. Pre-release preparations for juveniles may therefore require additional time or repeated exposure to novel foraging contexts. Encouragingly, these findings suggest that it is not “too late” for adult ‘alalā to incorporate new food resources – adult birds approached and contacted both novel and familiar fruit types at high rates, suggesting they may be suitable as release candidates.

Surprisingly, while birds were willing to contact both novel and familiar items, they were less likely to contact native fruits of any type compared to non-native options. This bias against native fruits, rather than explicit neophobia, may be more of a challenge to overcome in preparing individuals for release. While birds receive a rotation of both native and non-native fruits as enrichment, their main daily diet consists exclusively of non-native fruits (e.g., papaya, melon, apple). One possible explanation for the non-native fruit preference observed in this study is that even though the novel non-native fruits were unfamiliar, they nonetheless more closely resemble familiar items in their daily diet. There may also be other aspects of the appearance or content of native fruits that make these less preferred. For instance, while similar volumes of native and non-native fruit were provided, native fruits were individually smaller and berry-like, while large non-native fruits like watermelon were presented as cut pieces. Given that native fruits are a substantial component of the historic diet of wild ‘alalā (Sakai et al., 1986) and are foundational to their ecological role as seed dispersers (Culliney et al., 2012), this tendency to contact non-native food resources may need to be addressed to avoid hindering birds’ ability to integrate into native ecosystems.

Finally, while novelty did not impact patterns of contact, it did influence patterns of consumption. First, birds were overall less likely to consume test fruits than the control fruit, suggesting that degree of familiarity with a resource matters: even though birds had previously been exposed to the familiar test fruits, a highly familiar food item provided daily (papaya) was still more attractive. When comparing among the test fruits, we found an interaction between novelty and fruit type. Birds were similarly likely to consume non-native fruits regardless of novelty, but less likely to eat novel native fruits compared to familiar native items. Again, one possibility is that novel non-native fruits more closely resemble familiar foods in the birds’ daily diet, reducing dietary wariness. Another possibility is that there are salient differences in the intrinsic properties of native and non-native fruits that influence birds’ responses. For example, past work has shown that animals are more likely to consume a novel fruit when that item has a higher sugar content (Gosset & Roeder, 2001; Johnson, 2007; Visalberghi et al., 2003).

However, while comparative data on sugar content is limited for the fruits tested here, the gross energy content of the novel non-native fruit (dragonfruit, 3.78 kcal/g dry matter) and at least one of the novel native items (kolea, 4.16 kcal/g dry matter) appear roughly comparable (Puraikalan & Laton, 2023; Sakai & Carpenter, 1990).

From a management perspective, the specific hesitancy birds showed toward novel native foods suggests they may be reluctant to incorporate new fruit types into their diet post-release. One clear recommendation of this work is to reduce non-native fruits in the birds’ captive diet in favor of native items. Currently, however, sourcing sufficient quantities of native fruits for the daily diet of the conservation breeding population is not feasible. ‘Alalā identified for immediate release are exposed to native foods as part of the current pre-release process to ensure they will consume a minimum of several species prior to release. These efforts appear warranted and necessary given the preferences observed across the broader flock here. Expanding exposure to an even greater diversity of native species may further increase the likelihood that released birds will forage appropriately. More broadly, these results highlight a challenge inherent to long-term conservation breeding programs: behaviors important for survival in the wild may change over generations in human care, underscoring the importance of re-establishing wild populations as soon as conditions allow.

In addition, while this study focused on one key component of the historic diet of wild ‘alalā–native fruits–their diet included a broad range of items (including invertebrates, nectar, plant parts, eggs, nestlings, and small vertebrates) that merit further study. In particular, foraging for invertebrates made up a substantial portion of their historic feeding activity (55%, Sakai et al., 1986), and likely encouraged the evolution of their tool use (Rutz et al., 2016). Future research should also investigate which husbandry strategies most effectively facilitate contact and consumption of native items. Our finding that birds contacted novel native fruits at similar rates to familiar native fruits, but were less likely to consume them, aligns with past work indicating that dietary conservatism is more difficult to overcome than food neophobia (Marples et al., 2007). Successful strategies in other species have included extended exposure periods, repeated exposure, facilitated contact, and observational learning approaches (Greggor et al., 2016a; Gustafsson et al., 2014; Launchbaugh et al., 1997; Marples et al., 2007). A final consideration is that there may also be baseline variation in interest in different types of resources that is relevant for selection of birds for release. For example, while overall ‘alalā were least likely to consume novel native fruits, a few individual birds consumed *only* the novel native items.

When the reintroduction of a species depends entirely on individuals bred in human care, understanding how this population will respond to foraging and survival challenges in the wild is critical. This study reveals a preference for non-native fruits and wariness toward consuming novel native fruits in a conservation breeding population of the extinct-in-the-wild ‘alalā, two factors that may limit successful foraging in the wild if not addressed prior to release. While release sites are carefully selected with species-appropriate habitat and resources in mind, a preference for non-native foods may limit the foods that released birds incorporate into their diet, or lead birds into anthropogenically-modified areas. Understanding and mitigating behavioral biases will therefore be essential to ensure the success of future reintroductions.

## Acknowledgements

We would like to thank Kelsey Whittaker for her help in collecting data and to the animal care team at KBCC for helping facilitate this study. AC would like to acknowledge the support of a Doctoral Intern Fellowship from the Rackham Graduate School at the University of Michigan, which enabled this collaboration with the SDZWA.

## Author Contributions Statement

AG and BM conceptualized the study. AG coordinated data collection. AC analyzed the data and led the writing of the manuscript. All authors contributed critically to the drafts and gave final approval for publication.

## Notes

### Competing Interest Statement

The authors have declared no competing interest.

## References

Baldwin, P. H. (1969). The ‘Alalā (Corvus tropicus) of western Hawaii Island.’Elepaio, 30(5), 41–45.

Banko, P. C., Ball, D. L., & Banko, W. E. (2002). Hawaiian Crow (Corvus hawaiiensis). In A. Poole & F. B. Gill (Eds.), The Birds of North America (No. 648). Birds of North America, Inc., Philadelphia, PA.

Banko, P. C., Ball, D. L., & Banko, W. E. (2020). Hawaiian crow (Corvus hawaiiensis), version 1.0. In A. F. Poole & F. B. Gill (Eds.), Birds of the World. Cornell Lab of Ornithology.

Barrett, L. P., Flanagan, A. M., Masuda, B., & Swaisgood, R. R. (2024). The influence of pair duration on reproductive success in the monogamous ‘Alalā (Hawaiian crow, Corvus hawaiiensis). Frontiers in Conservation Science, 5, 1303239.

Bates, D., Mächler, M., Bolker, B., & Walker, S. (2015). Fitting linear mixed-effects models using lme4. Journal of statistical software, 67, 1–48.

Beck, B. B., Rapaport, L. G., Stanley□Price, M. R., & Wilson, A. C. (1994). Reintroduction of captive□born animals. In P. J. S. Olney, G. M. Mace, & A. T. C. Feistner (Eds.), Creative Conservation: Interactive Management of Wild and Captive Animals (pp. 265–286). Chapman & Hall.

Blum, C. R., Fitch, W. T., & Bugnyar, T. (2020). Rapid learning and long-term memory for dangerous humans in ravens (Corvus corax). Frontiers in Psychology, 11, 581794.

Bogale, B. A., Sugawara, S., Sakano, K., Tsuda, S., & Sugita, S. (2012). Long-term memory of color stimuli in the jungle crow (Corvus macrorhynchos). Animal Cognition, 15, 285–291.

Brown, C., & Laland, K. N. (2001). Social learning and life skills training for hatchery□reared fish. Journal of Fish Biology, 59(3), 471–493.

Brown, M. J., & Jones, D. N. (2016). Cautious crows: Neophobia in Torresian crows (Corvus orru) compared with three other corvoids in suburban Australia. Ethology, 122(9), 726–733.

Burnham, K.P., & Anderson, D.R. (2002). Model selection and multi-modal inference: A practical information-theoretic approach. Springer-Verlag.

Clarke, A. S., & Lindburg, D. G. (1993). Behavioral contrasts between male cynomolgus and lion□tailed macaques. American Journal of Primatology, 29(1), 49–59.

Crane, A. L., & Ferrari, M. C. (2017). Patterns of predator neophobia: a meta-analytic review. Proceedings of the Royal Society B: Biological Sciences, 284(1861), 20170583.

Culliney, S. B., Pejchar, L., Switzer, R. A., & Ruiz□Gutiérrez, V. (2012). Seed dispersal by a captive corvid: The role of the □Alalā (Corvus hawaiiensis) in shaping Hawai’i’s plant communities. Ecological Applications, 22(6), 1718–1732.

Ellis, D. H., Gee, G. F., Hereford, S. G., Olsen, G. H., Chisolm, T. D., Nicolich, J. M., … & Hatfield, J. S. (2000). Post-release survival of hand-reared and parent-reared Mississippi Sandhill Cranes. The Condor, 102(1), 104–112.

Ensminger, A. L., & Westneat, D. F. (2012). Individual and Sex Differences in Habituation and Neophobia in House Sparrows (Passer domesticus). ethology, 118(11), 1085–1095.

Glickman, S. E., & Sroges, R. W. (1966). Curiosity in zoo animals. Behaviour, 26(1–2), 151–188.

Gosset, D., & Roeder, J. J. (2001). Factors affecting feeding decisions in a group of black lemurs confronted with novel food. Primates, 42(3), 175–182.

Greenberg, R. S. (2003). The role of neophobia and neophilia in the development of innovative behaviour of birds. Animal innovation.

Greenberg, R., & Mettke□Hofmann, C. (2001). Ecological aspects of neophobia and neophilia in birds. Current Ornithology, 16, 119–178. (Springer).

Greggor, A. L., Thornton, A., & Clayton, N. S. (2015). Neophobia is not only avoidance: Improving neophobia tests by combining cognition and ecology. Current Opinion in Behavioral Sciences, 6, 82–89.

Greggor, A. L., McIvor, G. E., Clayton, N. S., & Thornton, A. (2016a). Contagious risk taking: social information and context influence wild jackdaws’ responses to novelty and risk. Scientific Reports, 6(1), 27764.

Greggor, A. L., Clayton, N. S., Fulford, A., & Thornton, A. (2016b). Street smart: Faster approach towards litter in urban areas by highly neophobic corvids and less fearful birds. Animal Behaviour, 117, 123–133.

Greggor, A. L., Vicino, G. A., Swaisgood, R. R., Fidgett, A., Brenner, D., Kinney, M. E., Farabaugh, S., Masuda, B., & Lamberski, N. (2018). Animal welfare in conservation breeding: Applications and challenges. Frontiers in Veterinary Science, 5, Article 323.

Greggor, A. L., Masuda, B., Flanagan, A. M., & Swaisgood, R. R. (2020). Age□related patterns of neophobia in an endangered island crow: Implications for conservation and natural history. Animal Behaviour, 160, 61–68.

Greggor, A. L., Sheppard, J., Masuda, B., Gaudioso-Levita, J., & Swaisgood, R. R. (2024). The influence of feeding station location on the space use and behavior of reintroduced ‘alalā: Causes and consequences. Conservation Science and Practice, 6(2), e13077.

Grodzinski, U., & Clayton, N. S. (2010). Problems faced by food-caching corvids and the evolution of cognitive solutions. Philosophical Transactions of the Royal Society B: Biological Sciences, 365(1542), 977–987.

Grueber, C. E., Nakagawa, S., Laws, R. J., & Jamieson, I. G. (2011). Multimodel inference in ecology and evolution: challenges and solutions. Journal of evolutionary biology, 24(4), 699–711.

Gustafsson, E., Saint Jalme, M., Bomsel, M. C., & Krief, S. (2014). Food neophobia and social learning opportunities in great apes. International Journal of Primatology, 35, 1037–1071.

IUCN. (2025). The IUCN Red List of Threatened Species. Version 2025-2.

Johnson, E. C. (2007). Rhesus macaques (Macaca mulatta) are not neophobic toward novel food with a high sugar content. American Journal of Primatology, 69(5), 591–596.

King, A. J., Fürtbauer, I., Mamuneas, D., James, C., & Manica, A. (2013). Sex-differences and temporal consistency in stickleback fish boldness. PLoS One, 8(12), e81116.

Kleiman, D. G. (1989). Reintroduction of captive mammals for conservation. BioScience, 39(3), 152–161.

Launchbaugh, K. L., Provenza, F. D., & Werkmeister, M. J. (1997). Overcoming food neophobia in domestic ruminants through addition of a familiar flavor and repeated exposure to novel foods. Applied Animal Behaviour Science, 54(4), 327–334.

Lenth, R., Singmann, H., Love, J., Buerkner, P., & Herve, M. (2018). Emmeans: Estimated marginal means.

Marples, N. M., & Kelly, D. J. (1999). Neophobia and dietary conservatism: two distinct processes?. Evolutionary Ecology, 13, 641–653.

Marples, N. M., Quinlan, M., Thomas, R. J., & Kelly, D. J. (2007). Deactivation of dietary wariness through experience of novel food. Behavioral Ecology, 18(5), 803–810.

Mathews, F., Orros, M., McLaren, G., Gelling, M., & Foster, R. (2005). Keeping fit on the ark: assessing the suitability of captive-bred animals for release. Biological Conservation, 121(4), 569–577.

Mettke□Hofmann, C., Winkler, H., & Leisler, B. (2002). The significance of ecological factors for exploration and neophobia in parrots. Ethology, 108(3), 249–272.

Mettke□Hofmann, C. (2014). Cognitive ecology: ecological factors, life□styles, and cognition. Cognitive Science, 5(3), 345–360.

Miller, R., Lambert, M. L., Frohnwieser, A., Brecht, K. F., Bugnyar, T., Crampton, I., Garcia□Pelegrin, E., Gould, K., Greggor, A. L., Izawa, E.-I., Kelly, D. M., Li, Z., Luo, Y., Luong, L. B., Massen, J. J. M., Nieder, A., Reber, S. A., Schiestl, M., Seguchi, A., Sepehri, P., Stevens, J. R., Taylor, A. H., Wang, L., Wolff, L. M., Zhang, Y., & Clayton, N. S. (2022). Socio-ecological correlates of neophobia in corvids. Current Biology, 32(1), 74–85.e4.

Miller, K. E., Hewett Ragheb, E. L., & Layman, C. A. (2024). Optimizing avian translocation success: A systematic review of the effect of release age on survival, dispersal, and productivity. Conservation Science and Practice, 6(9), e13195.

Owen-Smith, N., & Festa-Bianchet, M. (2003). Foraging behavior, habitat suitability, and translocation success, with special reference to large mammalian herbivores. Animal behavior and wildlife conservation, 93e109.

Puraikalan, Y., & Laton, A. (2023). Nutritional value of red-fleshed and skinned pitaya (Hylocereus polyrhizus) species. Int. J. Home Sci, 9(3), 271–275.

R Core Team. (2024) R: A Language and Environment for Statistical Computing. R Foundation for Statistical Computing.

Roberts, J. L., & Luther, D. (2023). An exploratory analysis of behavior-based and other management techniques to improve avian conservation translocations. Biological Conservation, 279, 109941.

Rutz, C., Klump, B. C., Komarczyk, L., Leighton, R., Kramer, J., Wischnewski, S., … & Masuda, B. M. (2016). Discovery of species-wide tool use in the Hawaiian crow. Nature, 537(7620), 403–407.

Sakai, H. F., Ralph, C. J., & Jenkins, C. D. (1986). Foraging ecology of the Hawaiian crow, an endangered generalist. The Condor, 88(2), 211–219.

Sakai, H. F., & Carpenter, J. R. (1990). The variety and nutritional value of foods consumed by Hawaiian crow nestlings, an endangered species. The Condor, 92(1), 220–228.

Shier, D. M. (2016). Manipulating animal behavior to ensure reintroduction success. In O. Berger-Tal & D. Saltz (Eds.), Conservation Behavior: Applying Behavioral Ecology to Wildlife Conservation and Management (pp. 275–304). Cambridge University Press.

Stoinski, T. S., Beck, B. B., Bloomsmith, M. A., & Maple, T. L. (2003). A behavioral comparison of captive-born, reintroduced golden lion tamarins and their wild-born offspring. Behaviour, 137–160.

Stoinski, T. S., & Beck, B. B. (2004). Changes in locomotor and foraging skills in captive□born, reintroduced golden lion tamarins (Leontopithecus rosalia rosalia). American Journal of Primatology, 62(1), 1–13.

Therneau, T. (2015). Mixed effects Cox models. CRAN repository.

U.S. Fish and Wildlife Service. (2009). Revised Recovery Plan for the ‘Alalā (Corvus hawaiiensis). Portland, Oregon.

Visalberghi, E., Sabbatini, G., Stammati, M., & Addessi, E. (2003). Preferences towards novel foods in Cebus apella: the role of nutrients and social influences. Physiology & Behavior, 80(2-3), 341–349.

Webster, S. J., & Lefebvre, L. (2000). Neophobia by the Lesser-Antillean Bullfinch, a foraging generalist, and the Bananaquit, a nectar specialist. The Wilson Bulletin, 112(3), 424–427.

