## Supplemental Materials for "Responses to novel food resources in the extinct-in-the-wild ‘alalā"

Latency to contact control fruit:

We analyzed latency to contact the control fruit (papaya) as a comparison for our main analyses of latency to contact the test fruit. We again included only birds who approached the testing setup (N = 185 trials). In contrast to the test fruit, we did not find a difference between juvenile and adult birds’ likelihood of contacting the control fruit (*β* = 0.07, *z* = 0.48, RI = 0.36; Table S2). We also found that birds contacted the control fruit less when it was paired with a non-native fruit compared to a native fruit (*β* = -0.87, *z* = 4.10, RI = 1.00; Table S2), but found no difference based on novelty (*β* = 0.05, *z* = 0.39, RI = 0.49; Table S2).

| (Intercept) | **Age Class** | **Fruit Category** | **Fruit Novelty** | **Sex** | **Category*Novelty** | **df** | **logLik** | **AICc** | **delta** | **weight** |
| --- | --- | --- | --- | --- | --- | --- | --- | --- | --- | --- |
| + |  | + |  |  |  | 1 | -621.59 | 1245.2 | 0.00 | 0.22 |
| + | + | + |  |  |  | 2 | -621.06 | 1246.2 | 0.99 | 0.14 |
| + |  | + | + |  |  | 2 | -621.13 | 1246.4 | 1.15 | 0.13 |
| + |  | + | + |  | + | 3 | -620.33 | 1246.8 | 1.62 | 0.10 |
| + |  | + |  | + |  | 2 | -621.39 | 1246.9 | 1.65 | 0.10 |
| + | + | + | + |  |  | 3 | -620.58 | 1247.4 | 2.14 | 0.08 |
| + | + | + | + |  | + | 4 | -619.80 | 1247.9 | 2.70 | 0.06 |
| + | + | + |  | + |  | 3 | -620.89 | 1248.0 | 2.75 | 0.06 |
| + |  | + | + | + |  | 3 | -620.94 | 1248.1 | 2.85 | 0.05 |
| + |  | + | + | + | + | 4 | -620.14 | 1248.6 | 3.38 | 0.04 |
| + | + | + | + | + |  | 4 | -620.43 | 1249.2 | 3.95 | 0.03 |

**Table S1.** Predictors included in averaged models of latency to contact control fruit.

|  | **Estimate** | **Std. Error** | **z value** | **Pr(>\|z\|)** | **RI** |
| --- | --- | --- | --- | --- | --- |
| **AgeClassAdult** | 0.07 | 0.14 | 0.48 | 0.63 | 0.36 |
| **FruitCategoryNonNative** | **-0.87** | **0.21** | **4.10** | **0.00** | **1.00** |
| **FruitNoveltyNovel** | 0.05 | 0.14 | 0.39 | 0.70 | 0.49 |
| **SexMale** | 0.03 | 0.14 | 0.21 | 0.83 | 0.28 |
| **Category*Novelty** | 0.10 | 0.25 | 0.39 | 0.69 | 0.20 |

**Table S2.** Output of averaged Cox proportional hazards model testing factors influencing the likelihood of contacting the control fruit.

Consumption of test fruit:

To ensure that our main findings regarding consumption of the test fruits were not driven by a lack of interest in eating during the trial, we repeated our analyses using only individuals who were observed eating at least one fruit (control, test, or both) during the trial (N = 123 trials). We found a similar, though slightly weaker, pattern to our main results, with an interaction between fruit category and fruit novelty suggesting that novelty reduces consumption of native but not non-native fruits (*β* = 1.71, *z* = 1.59, RI = 0.88; Table S4).

| (Intercept) | **Age Class** | **Fruit Category** | **Fruit Novelty** | **Sex** | **Category*Novelty** | **df** | **logLik** | **AICc** | **delta** | **weight** |
| --- | --- | --- | --- | --- | --- | --- | --- | --- | --- | --- |
| -0.40 |  | + | + |  | + | 5 | -66.85 | 144.2 | 0.00 | 0.44 |
| -0.64 | + | + | + |  | + | 6 | -66.55 | 145.8 | 1.61 | 0.20 |
| -0.56 |  | + | + | + | + | 6 | -66.66 | 146.0 | 1.83 | 0.18 |
| -0.75 |  | + | + |  |  | 4 | -69.23 | 146.8 | 2.60 | 0.12 |
| -0.77 | + | + | + | + | + | 7 | -66.39 | 147.7 | 3.54 | 0.07 |

**Table S3.** Predictors included in averaged models of test fruit consumption (by individuals who consume at least one fruit during the trial).

|  | **Estimate** | **Std. Error** | **Adjusted SE** | **z value** | **Pr(>\|z\|)** | **RI** |
| --- | --- | --- | --- | --- | --- | --- |
| (Intercept) | -0.54 | 0.46 | 0.47 | 1.16 | 0.25 |  |
| **FruitCategoryNonNative** | 0.19 | 0.70 | 0.71 | 0.27 | 0.79 | 1.00 |
| **FruitNoveltyNovel** | -1.71 | 0.66 | 0.67 | 2.55 | 0.01 | 1.00 |
| **CategoryNonNative***  **NoveltyNovel** | **1.71** | **1.07** | **1.07** | **1.59** | **0.11** | **0.88** |
| **AgeClassAdult** | 0.11 | 0.34 | 0.35 | 0.32 | 0.75 | 0.27 |
| **SexMale** | 0.07 | 0.27 | 0.27 | 0.26 | 0.79 | 0.25 |

**Table S4.** Output of averaged GLMM predicting likelihood of eating the test fruit (by individuals who consume at least one fruit during the trial).

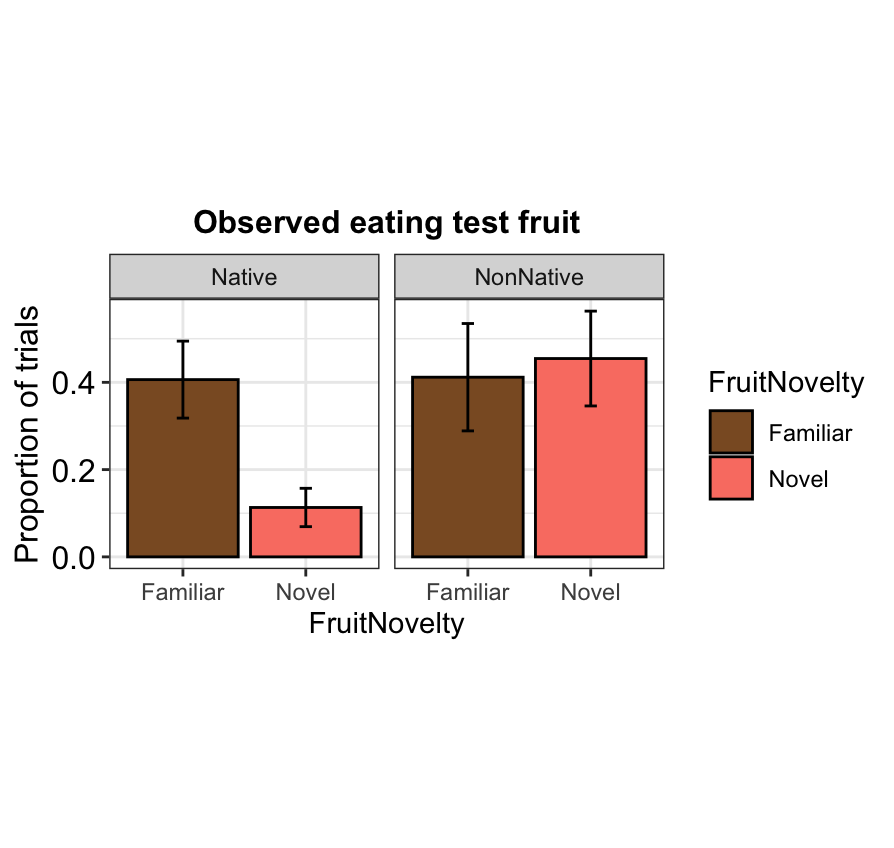

**Figure S1.** Patterns of test fruit consumption by novelty and category (by individuals who consume at least one fruit during the trial).
